# Centenarian Hotspots in Denmark

**DOI:** 10.1101/170654

**Authors:** Anne Vinkel Hansen, Laust Hvas Mortensen, Rudi Westendorp

## Abstract

**Background:** The study of regions with high prevalence of centenarians is motivated by a desire to find determinants of healthy ageing. While existing research has focused on selected candidate geographical regions, we explore the existence of hotspots in the whole the Denmark, which is a small and homogeneous country.

**Methods:** We performed a Kulldorff spatial scan across the whole of Denmark, searching for regions of birth, and regions of residence at age 71, where a significantly increased percentage of the cohort born 1906-1915 became centenarians. Next we compared mortality hazards for the identified regions to the rest of the country by sex and residence at age 71.

**Results:** We found a birth hotspot of 222 centenarians, 1.37 times more than the expected number, centered on a group of fairly remote rural islands. Significantly lower mortality hazards from age 71 onwards were confined to those who were born within the hotspot and persisted over a period of at least 30 years. At age 71, we found two residence-based hotspots of 348 respectively 238 centenarians, equaling 1.46- and 1.44 times the expected number. One is located in the high-income suburbs of the Danish capital and here the lower mortality hazard was confined to those who moved into the hotspot. In the second residence-based hotspot, both those who were born, and those who moved into the hotspot, showed significant lower mortality hazards.

**Conclusion:** Within the whole of Denmark, we identified several centenarian hotspots that have different biological underpinnings. These outcomes point to complex gene-environmental interactions explaining a variety of longevity trajectories.

## Introduction

Those who reach a hundred years capture our imagination. While the specific number is an arbitrary marker, becoming a centenarian is not a meaningless indicator of longevity, and the interest in geographical regions with high prevalence of centenarians has been ongoing at least since the start of the 20th century (Poulain et al 2013). The first of the current generation of well-validated longevity regions is the Sardinian “blue zone” (Poulain et al 2004), a small group of villages with a particularly high concentration of centenarians per 1000 births from 1880 to 1900. Other “blue zones” have been identified in the Okinawa region in Japan (Willcox et al 2008), the Ikaria island in Greece (Panagiotakos et al 2010) and the Nicoya peninsula in Costa Rica (Rosero-Bixby et al 2013). So far, the known longevity hotspots are in isolated, economically disadvantaged regions.

A plethora of explanatory factors has been proposed, roughly separating into seeing any effect as either genetic, rooted in local culture, or caused by local physical environment. For high-longevity zones in Calabria, Italy, areas with high prevalence of centenarians have been noted to also have a low variety of surnames, hinting at an explanation grounded in inbreeding (Montesanto et al 2007). Topographic factors, particularly altitude and steepness of terrain have been suggested (Pes et al. 2013). Life style factors have been investigated with varying results - the population of the Nicoya region have a diet rich in fibres, proteins and trans fats (Rosero-Bixby et al 2013), while Okinawa is the only blue zone to have a significantly lower caloric intake than the reference population (Willcox et al 2007). For the Nicoya blue zone, it has been observed that the effect is exclusive to those born and resident in the Nicoya region, with a non-significant decrease in mortality among immigrants and no decrease in mortality among out-migrants. This hints at the effect being, in some way, one of physical environment, or at least not tied to cultural factors carried along by out-migrants. As places change over time (sanitation improves, public transportation replaces walking, pesticide use changes), exposure to a place may mean different things in different decades. To our knowledge, there has not been any exploration of whether the effects observed in the various blue zones persist in generations after the ones they were detected in.

In terms of longevity, Denmark is a curious case. The country is small, socially homogeneous, highly economically developed, and has one of the world’s most generous universal welfare systems. At the same time, Denmark also has considerable social inequalities in health and a comparatively modest life expectancy, which is on par with that of the US. This study explores whether centenarian hotspots based on place of birth and place of residence late in life exist in Denmark. We explore the effects of time by examining whether such hotspots also prolong life in those moving there at a later age, and whether the effects remain in subsequent generations.

## Methods

We constructed a cohort of men and women born in Denmark from 1906 to 1915 and still alive at the age of 71 (after the introduction of the civil registration system in 1968). For each person, we identified parish of birth and place of residence by age 71 plus data on date of death and emigration. The data source was the national registry in which full follow-up for change of residence, death and emigration is possible for all Danish residents since 1968 (Pedersen 2011). Place of birth is recorded for all persons alive and resident in Denmark from 1977 and onwards. Information on later-life socioeconomic position and marital status was sourced from the 1970 census.

We searched for clusters of centenarians using a Kulldorf spatial scan (Kulldorf 1997) as implemented in the R package SpatialEpi. For each parish, the number of centenarians and the expected number based on number of births by sex and year of birth were assigned to the geographical centroid of the parish. The method then constructed zones as circular areas containing up to a pre-specified proportion (we chose to set this to 5 %) of the total population. For each zone, a likelihood ratio test statistic was constructed assuming a Poisson distribution of the number of observed centenarians. The zone most likely to be a cluster was selected as the one maximizing the test statistic, and significance measures were computed by Monte Carlo simulation. Any other zones for which the test statistic was significant at the pre-specified significance level (set to 0.05) were flagged as secondary clusters.

Having detected a hotspot, we compared mortality and centenarian proportion in people born inside and outside the hotspot for men and women. Mortality was compared via hazard ratios computed by Cox regression. We also compared mortality and centenarian proportion according to whether the subjects were born and stayed in, had left, were born outside and had moved to, or were born and had stayed in the cluster by age 71.

In order to explore the extent to which the effect was stable over time, we compared cohort mortality for those born inside and outside the cluster over the period from 1978 to 2015 for the cohorts with birth years 1906-1915, 1916-1925 and 1926-1935. Five-year mortality rate ratios were computed by Poisson regression.

As a sensitivity analysis, since a large percentage of observations were coded with municipality rather than parish of birth, we performed cluster detection where the observations coded with municipality were weighted out across the parishes whose centroids were inside that municipality. The weights were proportional to the size of the birth cohort by year and sex in each parish.

We also performed cluster detection based not on the proportion of a birth cohort reaching age 100, but on the proportion of those resident in a region by age 71 who become centenarians.

Analyses were done using R version 3.2.3 software.

## Results

There were 740,927 live births in Denmark from 1906 to 1915 (Statistics Denmark). Of these, we found 425,791 still alive and resident in Denmark by age 71. In order to make analyses by geographic location of birth, we made exclusions as follows: Those emigrating between age 71 and 100 (n = 1107), those lost in registry (n = 829), those with no parish of birth recorded (n = 57,178) and those missing information on parish of residence by age 71 (n=4,596). Of the group missing parish of birth, the majority (n = 42,436) had a record of municipality of birth in place of parish. Of the 362,064 individuals remaining in the study, 4,739 (1.3 %) reached the age of 100.

We first examined hotspots by parish of birth. This yielded one hotspot centered on a group of fairly remote rural islands (Langeland and encompassing rural areas on the islands of Funen and Lolland – see Figure 1). The birth hotspot had a total of 222 centenarians, 1.37 times the expected number, and the proportion was significantly larger than that of the remaining country with a p-value of 0.03. Per table 1, those born in the birth hotspot did not differ markedly from the baseline population on socioeconomic factors, except in being more likely to be homeowners (67 % vs 59 %).

**Figure 1:**
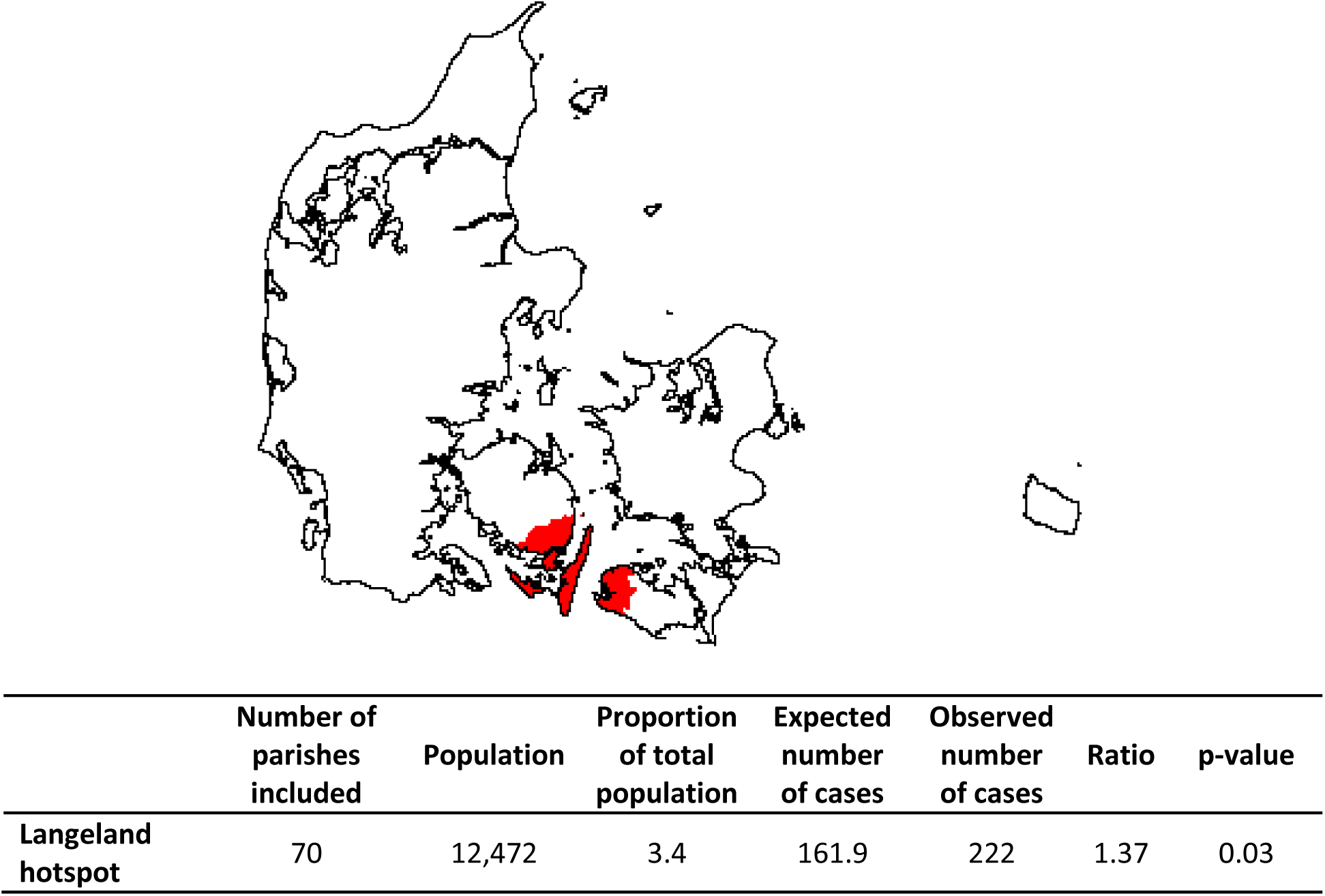
Hotspot for proportion of birth cohort surviving from 71 to 100.

**Table 1:** Socio-economic characteristics at age 71 for the study population and the birth-cohort centenarian hotspot

|  | Study population |  | Birth-cohort hotspot |  |
| --- | --- | --- | --- | --- |
|  | N | % | N | % |
| **Sex** |  |  |  |  |
| Female | 198,034 | 54.7 | 6,773 | 54.3 |
| Male | 164,030 | 45.3 | 5,699 | 45.7 |
| **Socioeconomic position in 1970** |  |  |  |  |
| 1 (highest) | 14,169 | 3.9 | 476 | 3.8 |
| 2 | 22,739 | 6.3 | 763 | 6.1 |
| 3 | 93,278 | 25.8 | 3,164 | 25.4 |
| 4 | 65,195 | 18 | 2,035 | 16.3 |
| 5 (lowest) | 89,038 | 24.6 | 3,285 | 26.3 |
| Pensioner | 70,348 | 19.4 | 2,471 | 19.8 |
| Other/unknown | 7,297 | 2 | 278 | 2.2 |
| **Marital status in 1970** |  |  |  |  |
| Unknown | 2,049 | 0.6 | 61 | 0.5 |
| Divorced | 18,750 | 5.2 | 547 | 4.4 |
| Married | 270,997 | 74.8 | 9,377 | 75.2 |
| Unmarried | 31,898 | 8.8 | 1,175 | 9.4 |
| Widow(er) | 38,370 | 10.6 | 1,312 | 10.5 |
| **Homeowner in 1970** |  |  |  |  |
| No | 147,632 | 40.8 | 4,156 | 33.3 |
| Yes | 214,432 | 59.2 | 8,316 | 66.7 |

Mortality differences between those born inside and outside the birth hotspot were more pronounced for women (HR 0.93, 95% CI 0.91 – 0.95) than for men (HR 0.97, 95% CI 0.95 – 1.00). This corresponds to the proportion surviving from 71 to 100 being 2.8 % and 1.9 % inside and outside the birth hotspot for women, and 0.6 % and 0.5 % for men (table 2).

**Table 2:** Mortality and centenarian proportion for those born in and outside the birth cohort centenarian hotspot, by sex and by place of residence at age 71. HR computed by Cox regression.

|  | N | Alive by age 100,<br>% (95 % CI) | HR (95% CI) | Adjusted*<br>HR (95% CI) |
| --- | --- | --- | --- | --- |
| **Hotspot** |  |  |  |  |
| Born outside hotspot | 349,592 | 1.3 (1.3 - 1.3) | 1 (REF) | 1 (REF) |
| Born in hotspot | 12,472 | 1.8 (1.6 - 2.0) | 0.95 (0.93 – 0.97) | 0.95 (0.94 – 0.97) |
| **Sex** |  |  |  |  |
| Men born outside hotspot | 158,331 | 0.5 (0.5 - 0.5) | 1 (REF) | 1 (REF) |
| Men born in hotspot | 5,699 | 0.6 (0.4 - 0.9) | 0.97 (0.95 – 1.00) | 0.98 (0.96 - 1.01) |
| Women born outside hotspot | 191,261 | 1.9 (1.9-2.0) | 1 (REF) | 1 (REF) |
| Women born in hotspot | 6,773 | 2.8 (2.4 – 3.2) | 0.93 (0.91 – 0.95) | 0.94 (0.91 – 0.96) |
| **Place of birth and residence at age 71** |  |  |  |  |
| Born and stayed outside hotspot | 346,230 | 1.3 (1.3 - 1.3) | 1 (REF) | 1 (REF) |
| Born outside and moved to hotspot | 3,362 | 1.3 (1 - 1.8) | 0.99 (0.96 – 1.03) | 1.01 (0.97 – 1.04) |
| Born in and left hotspot | 6,652 | 1.8 (1.5 - 2.2) | 0.96 (0.94 – 0.98) | 0.96 (0.93 – 0.98) |
| Born and stayed in hotspot | 5,820 | 1.8 (1.4 - 2.1) | 0.94 (0.92 – 0.97) | 0.95 (0.92 – 0.97) |

\*Adjusted for birth year, socioeconomic position, marital status and homeownership.

Mortality for those born in the birth hotspot and still remaining there by age 71 was slightly lower than for those who have left the birth hotspot (HR when compared to those born and remaining outside the hotspot 0.94, 95 % CI 0.92 - 0.97 and 0.96, 95 % CI 0.94 – 0.98 respectively) but with overlapping confidence intervals. There was no decrease in mortality for those born outside the birth hotspot but had moved into the birth hotspot by age 71 (HR 0.99, 95 % CI 0.96 – 1.03). The centenarian proportions reflect this pattern: the proportion surviving to 100 is 1.3% for those born outside and 1.8% for those born inside the birth hotspot, wherever they live by age 71.

Age-specific mortality rates for the cohorts born in the birth hotspot in the periods 1916-25 and 1926-35 are shown in the supplementary table. Age-specific mortality rates were comparable across birth cohorts, and remain significantly lower than expected from age 86 and onwards.

The sensitivity analysis including the group coded with municipality rather than parish of birth in the cluster detection process found the same primary hotspot, with 240 centenarian cases, 1.33 times the expected number with a p-value of 0.10.

When scanning for centenarian hotspots by place of residence at age 71 (see figure 2), we found one primary hotspot with 1.39 times (p-value 0.001), and one secondary hotspot with 1.33 times (p-value 0.003) the expected number of centenarians. The primary residence hotspot consisted of parishes in generally high-income suburbs in Northern Zealand. The secondary residence hotspot covered a region in Mid-Jutland. The total number of centenarians in each region was 379 and 337, respectively, for the primary and secondary residence hotspots. There were no further secondary clusters.

**Figure 2:**
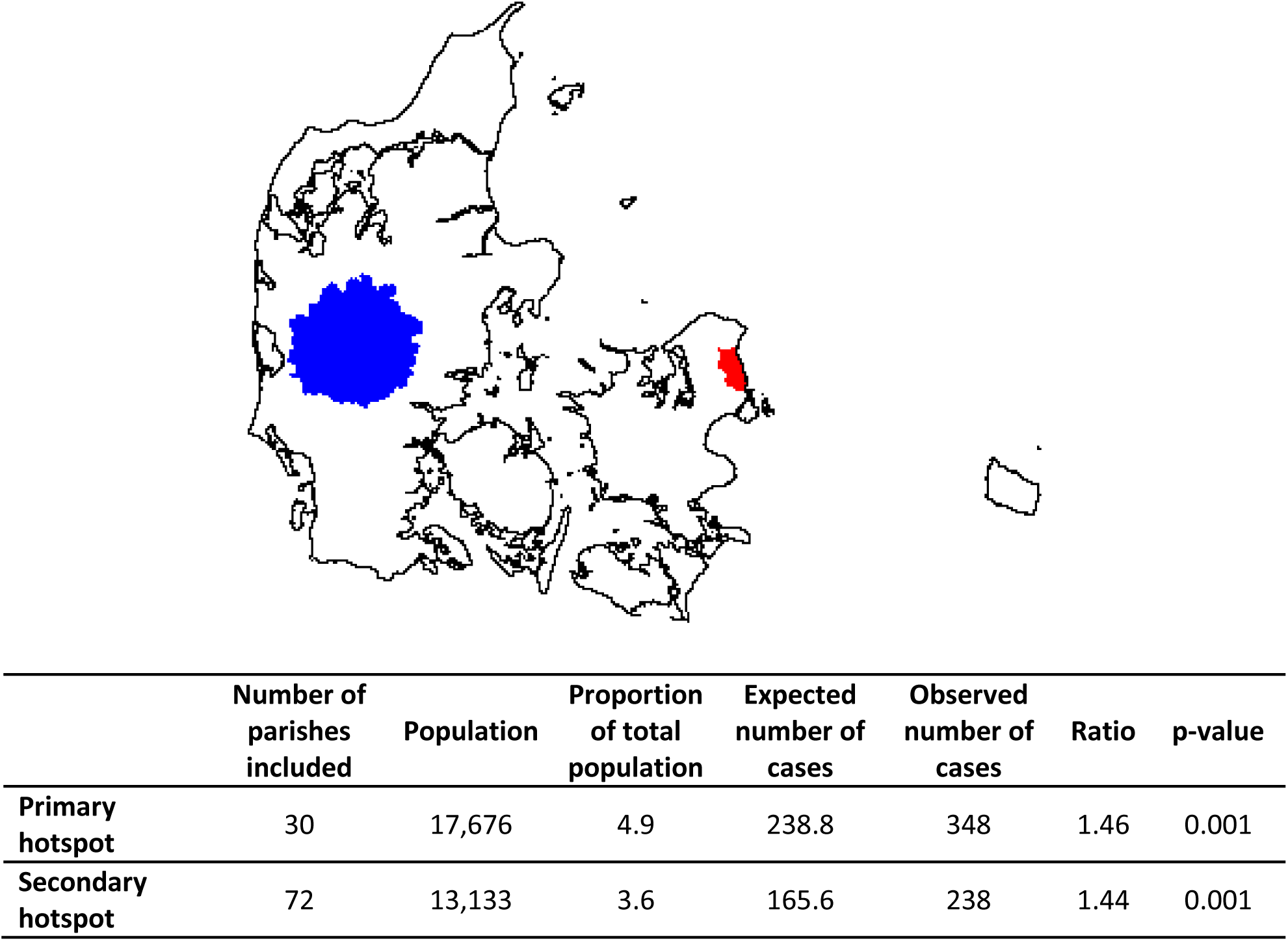
Primary (red) and secondary (blue) hotspots for proportion of residents in a region at age 71 surviving to age 100

Inhabitants of the primary residence hotspot in Northern Zealand showed a higher proportion with a high socioeconomic position (26 % in the two highest groups, compared to 10 % for the cohort in general) and a higher proportion of divorcees (Table 3) whereas no such differences were observed for the secondary residence hotspot in Mid-Jutland.

**Table 3:** Socio-economic characteristics at age 71 for the study population and the residence-based centenarian hotspots

|  | Study population |  | Primary residence-based hotspot |  | Secondary residence-based hotspot |  |
| --- | --- | --- | --- | --- | --- | --- |
|  | N | % | N | % | N | % |
| **Sex** |  |  |  |  |  |  |
| Female | 198,034 | 54.7 | 10,096 | 57.1 | 6,725 | 51.2 |
| Male | 164,030 | 45.3 | 7,580 | 42.9 | 6,408 | 48.8 |
| **Socioeconomic position in 1970** |  |  |  |  |  |  |
| 1 (highest) | 14,169 | 3.9 | 2,157 | 12.2 | 469 | 3.6 |
| 2 | 22,739 | 6.3 | 2,422 | 13.7 | 751 | 5.7 |
| 3 | 93,278 | 25.8 | 4,870 | 27.6 | 3,798 | 28.9 |
| 4 | 65,195 | 18.0 | 3,062 | 17.3 | 2,063 | 15.7 |
| 5 (lowest) | 89,038 | 24.6 | 2,433 | 13.8 | 3,376 | 25.7 |
| Pensioner | 70,348 | 19.4 | 2,311 | 13.1 | 2,441 | 18.6 |
| Other/unknown | 7,297 | 2 | 421 | 2.4 | 235 | 1.8 |
| **Marital status in 1970** |  |  |  |  |  |  |
| Divorced | 18,750 | 5.2 | 1,374 | 7.8 | 305 | 2.3 |
| Married | 270,997 | 74.8 | 12,999 | 73.5 | 10,323 | 78.6 |
| Unmarried | 31,898 | 8.8 | 1,403 | 7.9 | 1,282 | 9.8 |
| Widow(er) | 38,370 | 10.6 | 1,794 | 10.1 | 1,160 | 8.8 |
| Unknown | 2,049 | 0.6 | 106 | 0.6 | 63 | 0.5 |
| **Homeowner in 1970** |  |  |  |  |  |  |
| No | 147,632 | 40.8 | 9,186 | 52 | 2,483 | 18.9 |
| Yes | 214,432 | 59.2 | 8,490 | 48 | 10,650 | 81.1 |

Table 4 presents the matching mortality hazards for the residence-based hotspot from age 71 onwards. Within the primary hotspot, i.e. the high-income suburbs of the Danish capital, the significant lower mortality hazards were confined to those who moved into the hotspot (HR 0.88, 95 % CI 0.87 – 0.90), whereas those who were born in the same region did not show any benefit, irrespective whether they stayed or left the region (1.00, 95 % CI 0.96 – 1.04 and 1.00, 95 % CI 0.96 – 1.05 respectively). In the second residence-based hotspot, those who moved into the hotspot, showed significant lower mortality hazards (0.96, 95 % CI 0.93 – 0.98) but this was not different when compared to those who were born and stayed in the same region (0.95, 95 % CI 0.92 – 0.97).

**Table 4:** Mortality and centenarian proportion for those born in and outside the residence-based centenarian hotspots, by sex and by place of residence at age 71. HR computed by Cox regression.

|  | N | Alive by age<br>100,<br>% (95 % CI) | HR (95% CI) | Adjusted*<br>HR (95% CI) |
| --- | --- | --- | --- | --- |
| **Hotspots** |  |  |  |  |
| Resident outside hotspots | 331,255 | 1.3 (1.2 - 1.3) | 1 (REF) | 1 (REF) |
| Primary hotspot | 17,676 | 2.0 (1.8 - 2.2) | 0.89 (0.88 – 0.91) | 0.91 (0.89 – 0.93) |
| Secondary hotspot | 13,133 | 1.8 (1.6 - 2.1) | 0.95 (0.93 – 0.96) | 0.96 (0.95 – 0.98) |
| **Sex** |  |  |  |  |
| Men resident outside hotspots | 150,042 | 0.5 (0.4 - 0.5) | 1 (REF) | 1 (REF) |
| Men in primary hotspot | 7,580 | 0.8 (0.6 - 1.0) | 0.91 (0.89 – 0.93) | 0.94 (0.92 – 0.97) |
| Men in secondary hotspot | 6,408 | 0.9 (0.7 - 1.1) | 0.92 (0.90 – 0.94) | 0.94 (0.92 – 0.96) |
| Women resident outside hotspots | 181,213 | 1.9 (1.8 - 2.0) | 1 (REF) | 1 (REF) |
| Women in primary hotspot | 10,096 | 2.9 (2.6 - 3.2) | 0.90 (0.88 – 0.92) | 0.92 (0.90 – 0.94) |
| Women in secondary hotspot | 6,725 | 2.7 (2.3 - 3.1) | 0.94 (0.92 – 0.97) | 0.96 (0.94 – 0.99) |
| **Movement out of and into primary hotspot** |  |  |  |  |
| Born and stayed outside hotspot | 341,707 | 1.3 (1.2 - 1.3) | 1 (REF) | 1 (REF) |
| Born outside and moved to hotspot | 15,610 | 2.0 (1.8 - 2.3) | 0.88 (0.87 – 0.90) | 0.90 (0.89 – 0.92) |
| Born in and left hotspot | 2,681 | 1.1 (0.8 - 1.6) | 1.00 (0.96 – 1.04) | 1.00 (0.96 – 1.03) |
| Born and stayed in hotspot | 2,066 | 1.5 (1.1 - 2.1) | 1.00 (0.96 – 1.05) | 1.00 (0.96 – 1.05) |
| **Movement out of and into secondary hotspot** |  |  |  |  |
| Born and stayed outside hotspot | 340,103 | 1.3 (1.2 - 1.3) | 1 (REF) | 1 (REF) |
| Born outside and moved to hotspot | 5,473 | 2.0 (1.6 - 2.4) | 0.96 (0.93 – 0.98) | 0.97 (0.95 – 1.00) |
| Born in and left hotspot | 8,828 | 1.4 (1.2 - 1.7) | 0.95 (0.92 – 0.96) | 0.95 (0.93 – 0.97) |
| Born and stayed in hotspot | 7,660 | 1.7 (1.4 - 2.0) | 0.95 (0.92 – 0.97) | 0.96 (0.94 – 0.98) |

\*Adjusted for birth year, socioeconomic position, marital status and homeownership.

## Discussion

We found a Danish longevity birth hotspot centered on a group of fairly remote rural islands, with a 1.37 times increased chance of becoming a centenarian for the cohort born there 1906-1915. The effect is reflected in a lower post-71 mortality for both men and women, although the effect is markedly stronger in women. Mortality is lower for all those born in the hotspot, whether or not they have left by age 71. The difference in mortality is still observable and not substantially weakened for women born in the hotspot 1916-25 and 1926-35. We find two regions with significantly increased probabilities of reaching age 100 for those resident there at age 71, both with centenarian rates similar to those of the birth hotspot.

The main strength of this study is that the use of routinely collected, nation-wide registry data allows us to explore longevity distribution across an entire country for a 10-year birth cohort, that we have full follow-up from at least the age of 71, and that we can examine the effect in subsequent birth cohorts. One weakness is that we only have follow-up from 1977 and thus are unable to examine the extent to which our effects are shaped by mortality or emigration earlier in life. We only measure place of residence at two points in life, although people may have moved in and out of the regions studied. The Kulldorff cluster detection method is known to be conservative (Thomas et al 2003), and is constructed for point data, not area-aggregated data as we used. A likely effect of aggregation is over-estimation of cluster sizes (Jones & Kulldorff 2012), which, in turn, would reduce the estimate of the hotspot with centenarianism. The method allows for hotspots that group together regions separated by sea – this may seem counterintuitive, but arguably, a short stretch of sea is not necessarily a boundary in terms of local culture, population mixing or socioeconomic factors.

Nearly 10 % of observations were excluded due to missing information on parish of birth. When those with municipality of birth recorded were included, the location of the primary hotspot did not change, and while the effect size decreased past the point of insignificance, the absolute change was only from a ratio of 1.37 to a ratio of 1.33.

The absolute centenarian proportions in the study are not large. Comparison between studies are complicated by different outcome measures and especially different study periods, but the relative increase in probability of becoming a centenarian of 37 % can be compared to the 50 % increase in probability for the Sardinian blue zone or the three-fold increase in probability for the Sardinian restricted blue zone. Similarly, the age 70-100 death rate ratio of 0.95 for the Langeland hotspot compared to the rest of Denmark compares to the death rate ratio of 0.8 for Nicoyan men. Given the relatively high geographic, economic and cultural homogeneity in Denmark, it is not surprising that relative differences in centenarian proportions within Denmark are moderate.

Whatever the causes of the observed increases in extreme longevity for those born in birth hotspot, they seem to be determined before age 71. We see comparable post-71 mortality estimates for those born and staying in and leaving the hotspot, and no decrease in mortality for those moving to the birth hotspot as compared to those born and remaining outside the hotspot. The fact that there seems to be no benefits to moving to the birth hotspot later in life is reinforced by the results of the analysis of hotspots based residence at age 71 – the best place to be born is not the best place to grow old. This could point to the causes of the increase in centenarian prevalence being genetic or related to early life exposures, or being rooted in behaviors that are learned before old age and continued after leaving the region. The analyses of the subsequent cohorts suggest that whatever the determinants of longevity in the birth hotspot, they remain a factor over a period of at least 30 years.

When looking for causes for the birth hotspot, we should note that this area historically has been a poor area with an agricultural economy dominated by large estates. The main island of Langeland experienced mass emigration for a period leading up to the birth of the cohort studied - 30% of the population emigrated in the period 1868-1909 as compared to a national average of 10 % (Bender). While blue zones are generally poor and rural areas, this may be a rare case of those left behind by emigration being the healthier group, i.e. an ‘unhealthy migrant-effect’. Denmark is typically not considered a country with large genetic variation, but although the Danish population has historically been relatively mobile and there are no strong geographical barriers to mobility, there is still a tendency for surnames to cluster geographically, perhaps indicating lower mobility than might generally be the narrative (Boldsen 1992).

The interpretation of the residence at age 71 hotspots is less straightforward than that of the birth hotspot, given the nature of migration. Place of residence at age 71 is an indicator of socioeconomic and health status, and thus a straightforward interpretation of the secondary hotspots as “the best places to grow old” is not possible. They can just as well be interpreted as “the places where the fittest live by age 71”. Certainly, the Northern Zealand residential hotspot has markedly higher socioeconomic position by 1970 than the cohort in general and there could plausibly be selection factors into the Mid-Jutland hotspot as well.

## Supplementary material

**Table S1:**
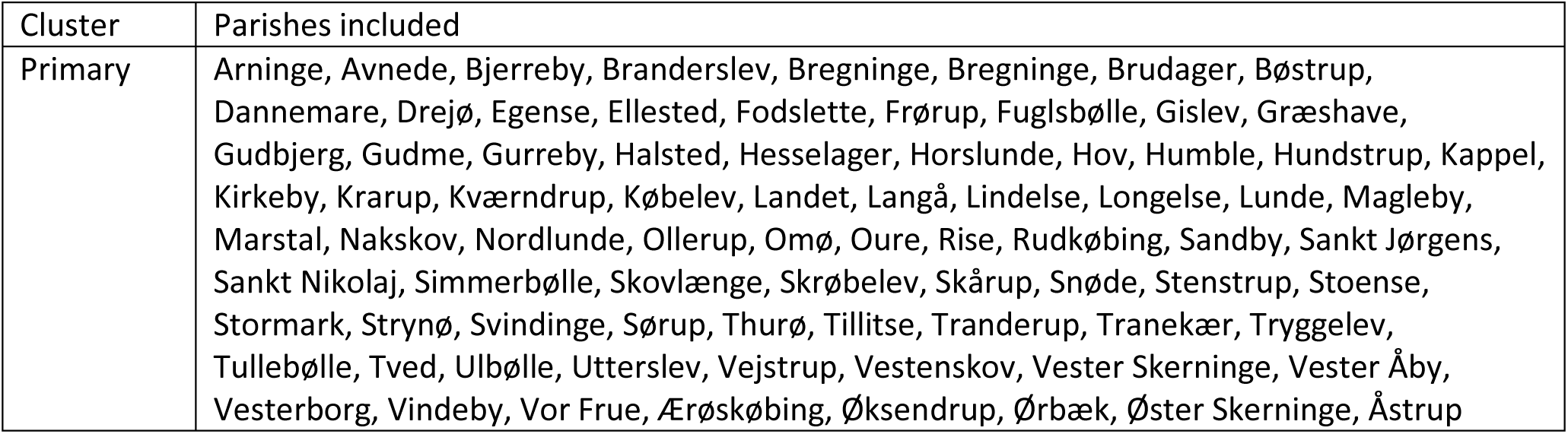
Parishes included in the birth cohort centenarian hotspot

**Table S2:**
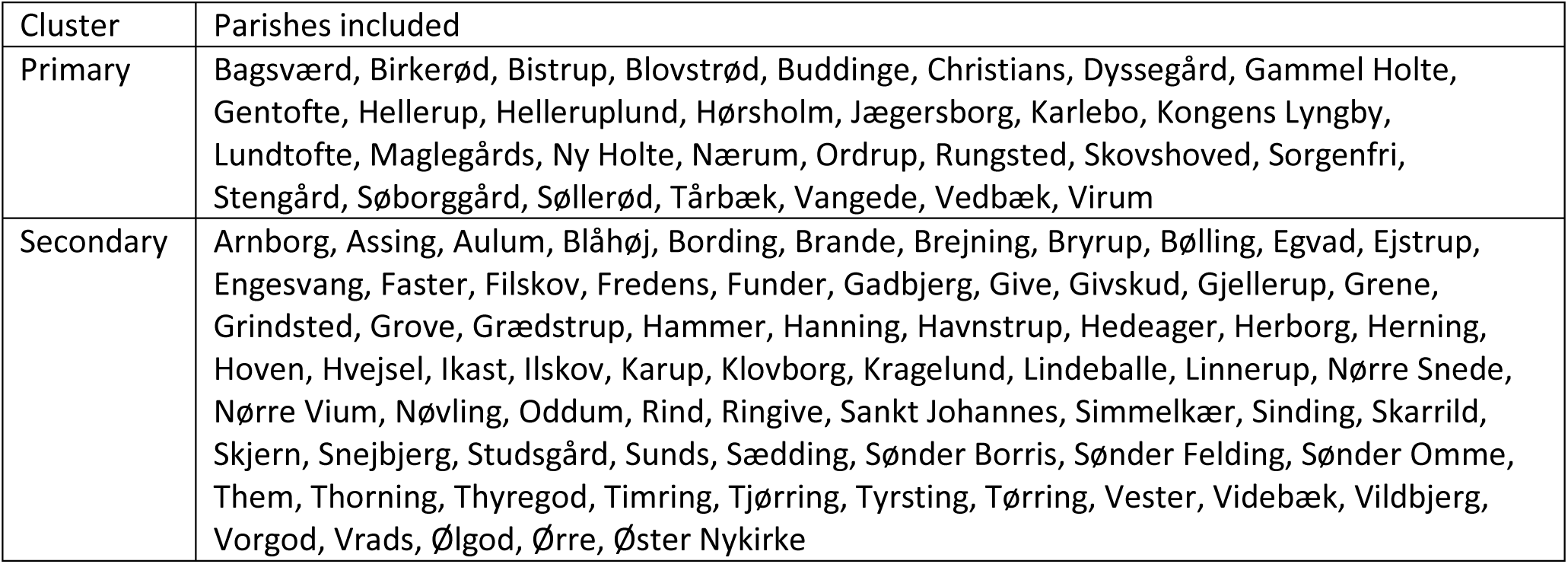
Parishes included in the primary and secondary residence-based centenarian hotspots

**Table S2:** Mortality rate ratios comparing the birth cohort centenarian hotspot to the rest of Denmark, for the cohorts born 1906-15, 1916-25 and 1926-35

|  | Birth year |  |  |
| --- | --- | --- | --- |
|  | 1906-1915 | 1916-1925 | 1926-1935 |
|  | Mortality rate ratio<br>(95 % CI) | Mortality rate ratio<br>(95 % CI) | Mortality rate ratio<br>(95 % CI) |
| **Population age 71, n** | 362,064 | 367,269 | 339,258 |
| **Born in hotspot, n** | 12,472 | 12,365 | 10,559 |
| **Age 71-75** | 0.98 (0.94 - 1.02) | 0.94 (0.90 - 0.98) | 1.00 (0.95 - 1.05) |
| **Age 76-80** | 0.98 (0.95 - 1.02) | 0.91 (0.88 - 0.95) | 0.98 (0.93 - 1.02) |
| **Age 81-85** | 0.94 (0.90 - 0.97) | 0.94 (0.91 - 0.98) |  |
| **Age 86-90** | 0.95 (0.91 - 0.98) | 0.94 (0.90 - 0.98) |  |
| **Age 90-95** | 0.92 (0.87 - 0.97) |  |  |
| **Age 96-100** | 0.91 (0.84 - 0.99) |  |  |

